# When to be a male? Role of resource-limitation and pollinators in determining floral sex in an andromonoecious spiderwort

**DOI:** 10.1101/2020.06.06.138354

**Authors:** Asawari Albal, G Azad, Saket Shrotri, Vinita Gowda

## Abstract

The evolution and maintenance of sexual systems in plants is often driven by resource allocation and pollinator preferences, and very little is known about their role in determining floral sex expression in plants. In annual, entomophilous plants three major constraints can be identified towards optimal reproduction: 1) nutrient resources available from the environment, 2) nutrient resources allocated towards reproduction, i.e., fruits vs. flowers, and 3) pollinator visitations.

Andromonoecy is a sexual system where plants bear both staminate and hermaphrodite flowers on the same inflorescence. The optimal resource allocation hypothesis suggests that under nutrient constraints, plants will produce more male flowers since they are energetically cheaper to produce over the more expensive hermaphrodite flowers. We test this hypothesis in the andromonoecious *Murdannia simplex* (Commelinaceae) by quantifying male and hermaphrodite flowers in a natural population and contrasting the distribution of the two sexes in plants from two resource conditions (stream population vs. plateau population). We next carried out choice experiments to test pollinator preference towards a specific sex.

We found that in *M. simplex*, production of hermaphrodite flowers is resource-dependent and under resource constraints fewer numbers of flowers were produced and most of them were males. We failed to observe pollinator preference towards either sex but *Amegilla spp*. and *Apis cerana* showed higher visitation towards the most abundant sex within a trial, suggesting frequency-dependent visitation. Thus, we conclude that environmentally driven resource constraints play a bigger role in driving floral sex expression in *Murdannia* over direct pollinator-driven constraints.

## Introduction

Andromonoecy refers to the sexual system where both staminate (male) flowers and perfect (hermaphrodite) flowers are present on the same plant (Bertin 1982a, Miller and Diggle 2003, Vallejo-Marín and Rausher 2007a, 2007b). In andromonoecious plants, the hermaphrodite flower fulfills the female function in the absence of a true ‘female’ flower, and it has been suggested that andromonoecy probably evolved from hermaphroditism by the loss of female reproductive structure (Lloyd 1980, Bertin 1982a). One of the most widely accepted hypotheses on the evolution of andromonoecy is the optimal resource allocation hypothesis which suggests that under resource limitation, male flowers will be produced instead of hermaphrodite flowers. This hypothesis rests upon the premise that male flowers are energetically cheaper (or economical) to produce because reproductive allocation towards them ends with pollen production (Anderson and Symon 1989, Narbona et al. 2002, Verdú et al. 2007), while females are expensive because fruit development and seed germination are energy-intensive physiological processes (Bertin 1982a, Kaul et al. 2002, Obeso 2002, Verdú et al. 2007).

Examples of reallocation of resources to male flowers (i.e. male bias) have been shown in several plant species where staminate flowers are produced when nutrient resources become scarce due to its utilization by the costlier gynoecia towards fruit production (Bertin 1982a, 1982b, May and Eugene Spears Jr 1988, Miller and Diggle 2003, Venkatesan 2017). A well-studied species is *Solanum hirtum* (Solanaceae) in which it has been shown that as a result of successful pollination, as the ovary develops, resources are relocated within an inflorescence to the male flowers, thus influencing the sex of successive flowers and their distribution (Diggle 1993, Miller and Diggle 2003). Another example comes from *Raphanus raphanistrum*, where Stanton et al. (1987) showed that in plants which were heavily pollinated, the number of ovules per flower decreased while unpollinated plants did not show any significant decrease in number of ovules. This suggests that plants can modify their reproductive output to adjust for the resource limitations faced by them.

In plants, resource limitation or nutrient limitation that can affect sex of flowers can be identified as: a) environmental nutrient limitation (Charnov and Bull 1977, Primack and Lloyd 1980a, Stephenson 1981, Bertin 1982a, Solomon 1985, Diggle 1993), and/or b) within-plant nutrient limitation (Stephenson 1981, May and Eugene Spears Jr 1988, Diggle 1993, Miller and Diggle 2003, Ortiz et al. 2003, Vallejo-Marín and Rausher 2007a). In *Aesculus californica, A. pavia, Leptospermum scoparium* and *Passiflora incarnata*, phenotypic sex expression (hermaphroditic inflorescences or hermaphroditic flowers) has been shown to vary as a response to environmental conditions or status of the nutrient resources, especially during fruit development when resources are limited (Benseler 1975, Primack and Lloyd 1980b, May and Eugene Spears Jr 1988). Further, it has been shown that the amount of resources that a plant acquires in the proximal/distal ends of the inflorescence can also vary. The proximal/ basal end of the inflorescence has more resources than its distal end, which results in more hermaphrodite flowers to be present at basal ranks of an inflorescence, while male flowers are relegated to the distal end (Diggle 1994, 1997, Lewis and Gibbs 1999, Ashman and Hitchens 2000, Miller and Diggle 2003, Kaul and Koul 2008).

Bateman’s principle (Wilson et al. 1994) asserts that-“Fitness gain through male function is limited primarily by mating opportunity, while fitness gain through female function is limited primarily by resource availability for offspring production”. A similar idea was proposed in the sex allocation theory which predicted that attracting pollinators that promote outcrossing by improving floral attractiveness (e.g., large petals, nectar availability) results in a higher fitness gain through the male function than female function (Elle and Meagher 2000). This is because male success is correlated with mating opportunities, while one or a few visits by pollinators can adequately fertilize all ovules. To explain the role of male flowers in andromonoecious plants, two not-mutually exclusive hypotheses have been proposed (Zhang and Tan 2009): i) Pollen donor hypothesis which emphasizes the function and reproductive success of male flowers. It predicts that in a resource-limited condition the wastage of nutrient resources by investment in the female reproductive structure is reduced by the production of only male flowers. ii) Pollinator attractor hypothesis which emphasizes the function and reproductive success of female flowers. It predicts that male flowers increase the floral display and attract more pollinators at a lower cost thereby increasing female reproductive success (of the hermaphrodite flowers) at lower resource investment.

In entomophilous plants, pollinator dependence and pollination success is often driven by floral characteristics (Armbruster 2001, Fenster et al. 2004).. Other studies have also shown that several characteristics of an inflorescence such as number of flowers, size of flowers, and reward of flowers play a critical role in attracting pollinators to the plant (Ortiz et al. 2003, Christopher et al. 2019). All of the above examples are from hermaphroditic reproductive systems, and the role of specific floral sex, and its contribution to reproductive output in an andromonoecious systems are not widely studied (although see Diggle 1993, Ashman et al. 2000, Ashman and Morgan 2004).

Previous studies in sexually dimorphic plants have shown that pollinators typically show a preference for the functionally male flowers over the functionally female flowers owing to different floral traits like petal length (Ashman et al. 2000), levels of pollen production (Biezychudek 1987, Eckhart 1991) and/or floral scents (Ashman et al. 2005), where these preferences are learned through foraging experience (Cresswell and Galen 1991). Zhang and Tan (Zhang and Tan 2009) were the first to examine the function of male flowers in the andromonoecious *Capparis spinosa* where they showed that the pollinators did not discriminate between the two sexes, despite morphological dimorphism. However, similar studies investigating sex-specific pollinator preferences and more specifically sex-specific pollinator preferences in sexually dimorphic plants that have near-identical morphological features are not known.

In this study, we explore the effect of nutrient resources as well as pollinator preference in determining the sex of flowers in a wild population of the andromonoecious *Murdannia simplex* (Commelinaceae, Spiderworts). The pantropical plant family Commelinaceae (Spiderworts) is known to have several genera that display andromonoecy. Despite the dominance of andromonoecy in this family, to date, most pollination studies within this family have focused only on identifying pollinators (Kaul et al. 2002, Williams and Walker 2003, Oziegbe et al. 2013, Sigrist and Sazima 2015), and only one study has explored the role of floral guides in pollinator attraction (Ushimaru et al. 2007). Based on predictions from the optimal resource allocation hypothesis and pollinator attractor hypothesis we address the following questions:

A. Does the proportion of male and hermaphroditic flowers in a population represent resource limitation?
B. Do pollinators show preference to male or hermaphrodite flowers?
C. Do male flowers improve floral display to maintain visitation rate and function as a cheaper source of outcrossing pollen?

## Materials and methods

*Murdannia simplex* (Vahl) Brenan (Commelinaceae) is a mass-flowering, sub-erect, annual, andromonoecious herb which is widely distributed in moist deciduous forests and grasslands of the Western Ghats of India (Nandikar and Gurav 2015; Fig. 1a). The plant is seasonal, 40-65 cm tall, and the shoot emerges once monsoonal rains begin. Flowering lasts from September to November (Nandikar and Gurav 2015) and flowers are borne on a cymose inflorescence. The inflorescence bears 5-20 purple-colored, three-petaled male and hermaphrodite flowers, and each flower measures ∼15-25 mm in diameter. The hermaphrodite flowers are characterized by the presence of a lateral pistil, with two upper fertile stamens, curved downwards and a third lower sterile stamen. Filaments have long purple, bearded hair. The flower also bears three trilobed staminodes with glabrous filaments. The male flowers are characterized only by the lack of a pistil (Fig. 1b - f). The flowers are defined by a short floral anthesis time, from 12:30 hrs. to 16:00 hrs., with the principal pollinator reward being pollen grains only (Faden 2000, Nandikar and Gurav 2015).

**Fig. 1.**
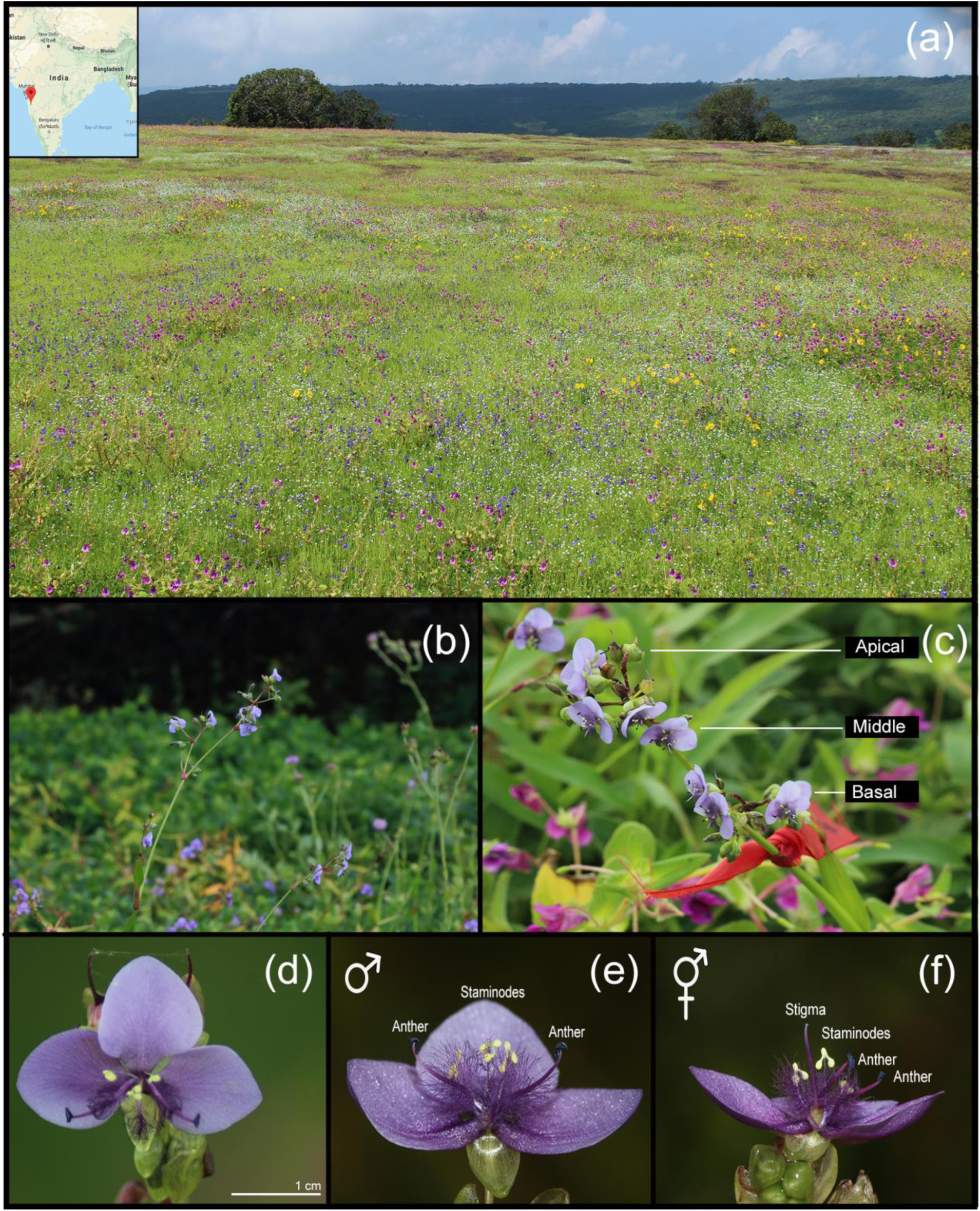
Habitat and floral features of *Murdannia simplex* (a) habitat on Kaas Plateau, Maharashtra during flowering season (Inset: Location in India), (b) *M. simplex* habit, (c) inflorescence image showing positions identified from base to the apex, (d) *M. simplex* flower front view, (e) *M. simplex* male flower, (f) *M. simplex* hermaphrodite flower. Photo courtesy: (a), (c), (d) - Asawari Albal, (b), (e), (f) - Saket Shrotri.

### Study site

Western Ghats is one of the four biodiversity ‘hotspots’ in India. The northern part of Western Ghats is famous for high elevation laterite plateaus and slopes, one of them being Kaas plateau (17° 43′ 12″ N, 73° 49′ 22″ E), which is located at an altitude of ∼1225 m above sea level near the city of Satara, Maharashtra and covers an area of ∼10 km^2^ (Fig. 1a). Kaas plateau was declared as a UNESCO World Heritage site in 2012 due to its highly endemic floral diversity, which is most abundant in the months of August to November.

Capped with red lateritic crusts, the upper Kaas Plateau is an arid habitat except during the monsoon season (Fig. 1a). It receives over 2500 mm of rain every year, mainly during the monsoon months (June to September), and the daily mean temperatures are over 22°C. Floral diversity on the Kaas plateau is a model representation of flora that is associated with seasonal monsoonal rains in the Western Ghats, as well as the flora of lateritic plateaus of the Western Ghats. All field experiments and population-level sex ratio measurements were carried out across three consecutive years - 2017 to 2019, between 15^th^ September and 5^th^ November. The first two years focused on the study of plant-pollinator interactions and studies on the effect of nutrient resources on sex of the flowers was carried out in the third year.

### Population-level sex ratios

To determine the extent of sexual dimorphism we compared flower sizes of the two sexes. Pairs of male and hermaphrodite flowers were photographed from 15 random inflorescences on a laminated graph sheet, and their petal length and width were measured manually (Online Resource 1: Fig. 1).

In order to identify the distribution pattern (ratios) of the two sexes within a plant and within an inflorescence, a multi-year (2018 and 2019) census was conducted. All census were carried out at an interval of seven days, across 4 weeks (spanning over one and a half months) covering the peak flowering season of *M. simplex*. Floral sex was determined between 12:30 hrs. to 16:00 hrs. (anthesis time of *M. simplex* flowers), on marked inflorescences (N = 30). Since a single individual can bear 2-5 inflorescences, a pooled census was taken from all the inflorescences of a plant. To identify if there was a positional-bias in the floral sex i.e. from basal to apical position within an inflorescence (see Stephenson 1981, Diggle 1997, Cuevas and Polito 2004), the positions of each flower and its sex was recorded throughout the census duratio. The positions within an inflorescence were divided into three categories-apical, middle, and basal as shown in Fig. 1c.

To test the effect of nutrient resources on floral sex expression, we counted male and hermaphrodite flowers in two populations that differed in the availability of water and other resources (presence of a stream and a deep soil substrate), which we predicted to be primary limiting resources on an otherwise homogeneous plateau. The first population is henceforth referred to as the plateau population - ‘Plateau_pop’, while the second population is referred to as the stream population - ‘Stream_pop’. Plateau_pop was selected as the resource-poor population whereas Stream_pop was selected as the resource-rich population. The two locations were separated by ∼576 meters and other than the presence of a stream and a deep soil substrate, can be viewed as a single population. The census was carried out in October 2019 on randomly selected individuals within these two populations (N = 100 per location).

### Pollination biology

Pollen is the only pollinator reward in the genus *Murdannia* (Faden 2000, Nandikar and Gurav 2015). To identify any quantifiable difference in the pollen or paternal contribution between the male and hermaphroditic flowers, the amount of pollen per anther in virgin flowers of *M. simplex* was quantified using a hemocytometer.

Pollinator visitations on *M. simplex* were quantified by observing pollinators in 2 ft. x 2 ft. observation arenas (N = 49) in multiple intervals of 15 minutes between 12:30 hrs. to 16:00 hrs. (anthesis time of *M. simplex*). Each arena consisted of 4-10 inflorescences with a total of 15-30 flowers. Visitation by a pollinator was recorded when a pollinator landed on the flower and actively collected pollen. In each observation interval, we recorded the total number of flowers present, type of pollinator, and number of pollinator visits. Pollinator visitation rate was then calculated as the total number of visits per flower per hour. At the end of the flowering season, fruitset was quantified by counting the total number of fruits in randomly selected inflorescences from the study population (N = 26 in 2017 and N = 40 in 2019).

### Sex-specific pollinator preference experiment

Manipulated choice experiments were designed and conducted in the wild to test a) presence of visitation bias by pollinators towards male flowers (M) and/or hermaphrodite flowers (H), and b) presence of floral-density bias by pollinators. It was hypothesized that if the pollinator has an inherent preference for any sex, the density of a specific sex in our experimental set up will not affect the pollinator’s visitation rate. Floral distributions within an inflorescence were defined to have binary outcomes i.e., male abundant or hermaphrodite abundant, or equally male and hermaphroditic. The experimental setup consisted of two pots that were kept one foot apart with a total of 5-15 flowers in each pot and pollinator visits were quantified by eye between 12:30 hrs. to 16:00 hrs. Three sex-specific floral-density treatments were designed: H < M (T1), H = M (T2) and H > M (T3) were tested. The density of flowers of the two sexes in each experimental set up was manually controlled and each treatment was repeated for a total of eight trials and a total observation time of ∼21.5 hours (Online Resource 1: Fig. 2). Unmanipulated, wild set up observations served as the control. The mean visitation rate per flower to male flowers and hermaphrodite flowers were calculated for each pollinator type, by dividing the total number of visits by a specific pollinator with the total number of flowers present in the experimental set up. Pollinator visitation rate was calculated as the total number of visits per flower per hour.

**Fig. 2.**
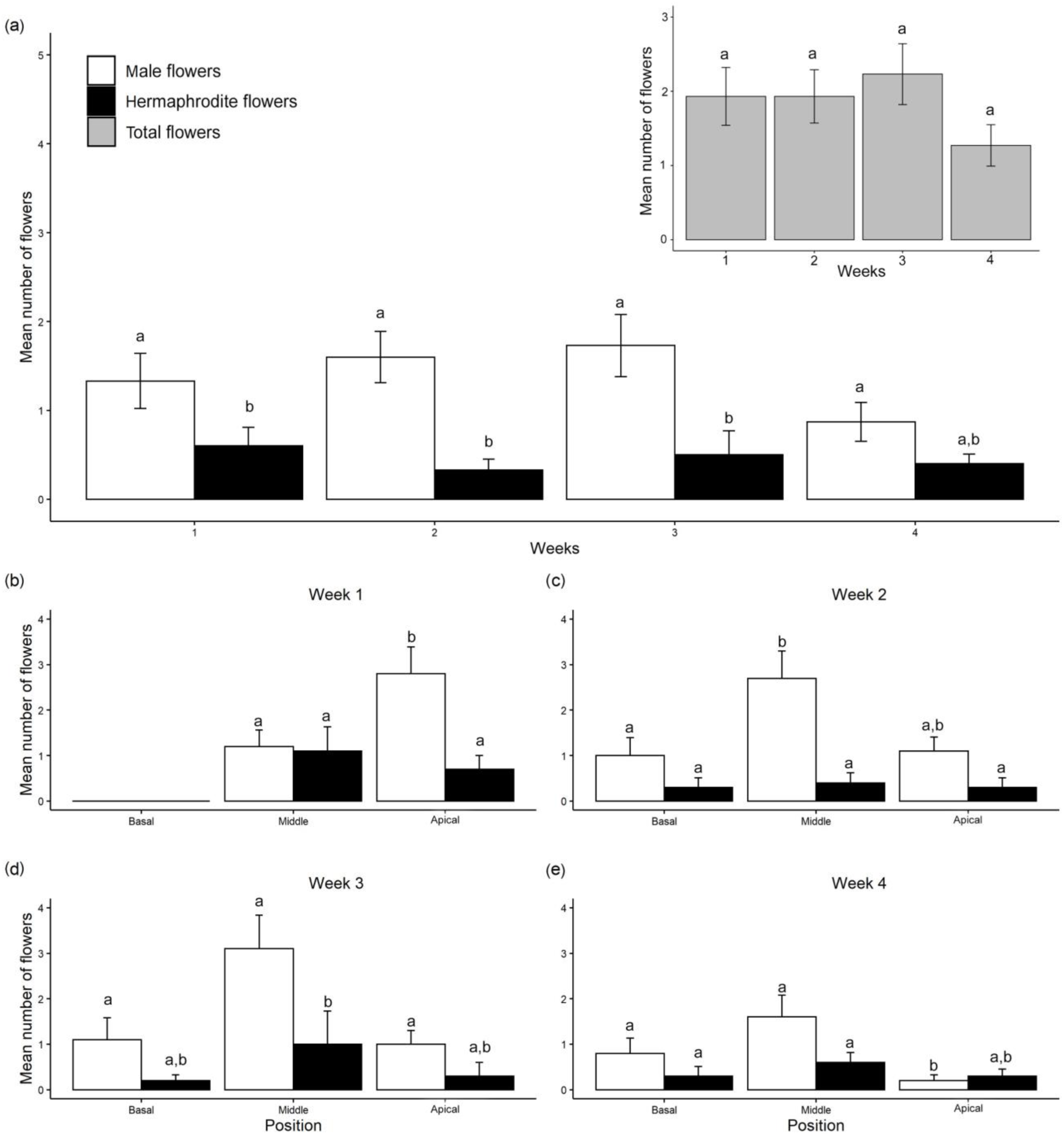
Variation in population-level sex ratios in *M. simplex* (a) by week from first flowering date (Kruskal-Wallis chi-squared = 38.7067, df = 7, *p*-value = 0.00; P<0.025) (Inset: Week-wise proportion of total number of flowers; Kruskal-Wallis chi-squared = 3.2847, df = 3, *p*-value = 0.3498, N = 30; P<0.025) (See Online Resource 1: Table 1) and (b) - (e) within-inflorescence from Week 1 - Week 4 respectively (Kruskal-Wallis chi-square test; df = 5, P<0.025). For (a) columns to be compared within the week and within sexes (Mean±SE). For (b) columns to be compared within the sexes and within positions, across all 4 weeks (Mean±SE). Significant *p*-values depicted by different alphabets (Online Resource 1: Table 3; P<0.025).

The visitation rates in the control treatments (natural visitations) were calculated and compared with the visitation rates obtained from the choice experiments (H < M, H = M and H > M). The mean visitation rates across all three treatments were also compared with the total number of flowers present in the trial to identify any effect of floral display size.

### Statistical analysis

All analyses were performed in R (R Core Team 2019) and raw data was tested for normality before proceeding with statistical analysis using Shapiro-Wilk’s method. When Gaussian type distribution could not be assumed, non-parametric Kruskal-Wallis rank-sum tests were run using the dplyr package (Wickham et al. 2020). The floral diameters and pollen densities per flower between male and hermaphrodite flowers were compared using unpaired two-sample t-test. The variation in proportion of male and hermaphrodite flowers was quantified using Kruskal-Wallis test, and post hoc Dunn’s test (Dinno 2017) was used to identify proportional difference between the two sexes across the four weeks. Simple linear regression was used to check the effect of mean number of male flowers, hermaphrodite flowers and total flowers on the fruit set respectively, and the effect of total floral display size on the mean visitation rate across the three treatments in the choice experiments. The alpha was set at 0.5 (P<0.025) for all statistical analysis. All graphical plots were produced using the ggplot2 package in R (Wickham 2016).

## Results

### Population-level sex ratios

Due to the small size and fragile nature of the flower, floral size was measured as the widths of the flowers (widest diameter). There was no significant difference between floral size of the male and hermaphrodite flowers (t = 0.11072, df = 28, *p*-value = 0.9126; Online Resource 1: Fig. 1). Throughout the flowering season, male flowers were present in higher proportions than the hermaphrodite flowers in all the study plots by at least 70-80% (Kruskal-Wallis chi-squared = 38.707, df = 7, *p*-value = 2.223^e-06^; Fig. 2a; Online Resource 1: Table 1). A slight decrease in the proportion of male flowers was observed only in week 4 (Mean±SE: Week 1 = 1.33±0.31, Week 2 = 1.6±0.29, Week 3 = 1.73±0.35, Week 4 = 0.87±0.22; Fig. 2a; Online Resource 1: Table 1).

In contrast, the number of hermaphrodite flowers remained similar across all four weeks of observation (Mean±SE: Week 1 = 0.60±0.21, Week 2 = 0.33±0.12, Week 3 = 0.50±0.27, Week 4 = 0.40±0.22 ; Fig. 2a; Online Resource 1: Table 1) and only plants that were closer to the water source i.e. stream population, had a significantly higher number of hermaphrodite flowers (Mean±SE: Plateau_pop H = 0.43±0.07 vs. Stream_pop H = 1.75±0.14; *p*-values = 0.0000; Fig. 3; Online Resource 1: Table 2). Overall, the mean number of total flowers remained consistent for the first two weeks (Mean±SE: Week 1 = 1.93±0.39, Week 2 = 1.93±0.37 flowers; Fig. 2a inset), increased in week 3 (2.23±0.41 flowers) and then reduced in week 4 (1.26±0.28 flowers; Kruskal-Wallis chi-squared = 3.2847, df = 3, *p*-value = 0.3498; Online Resource 1: Table 1).

**Fig. 3.**
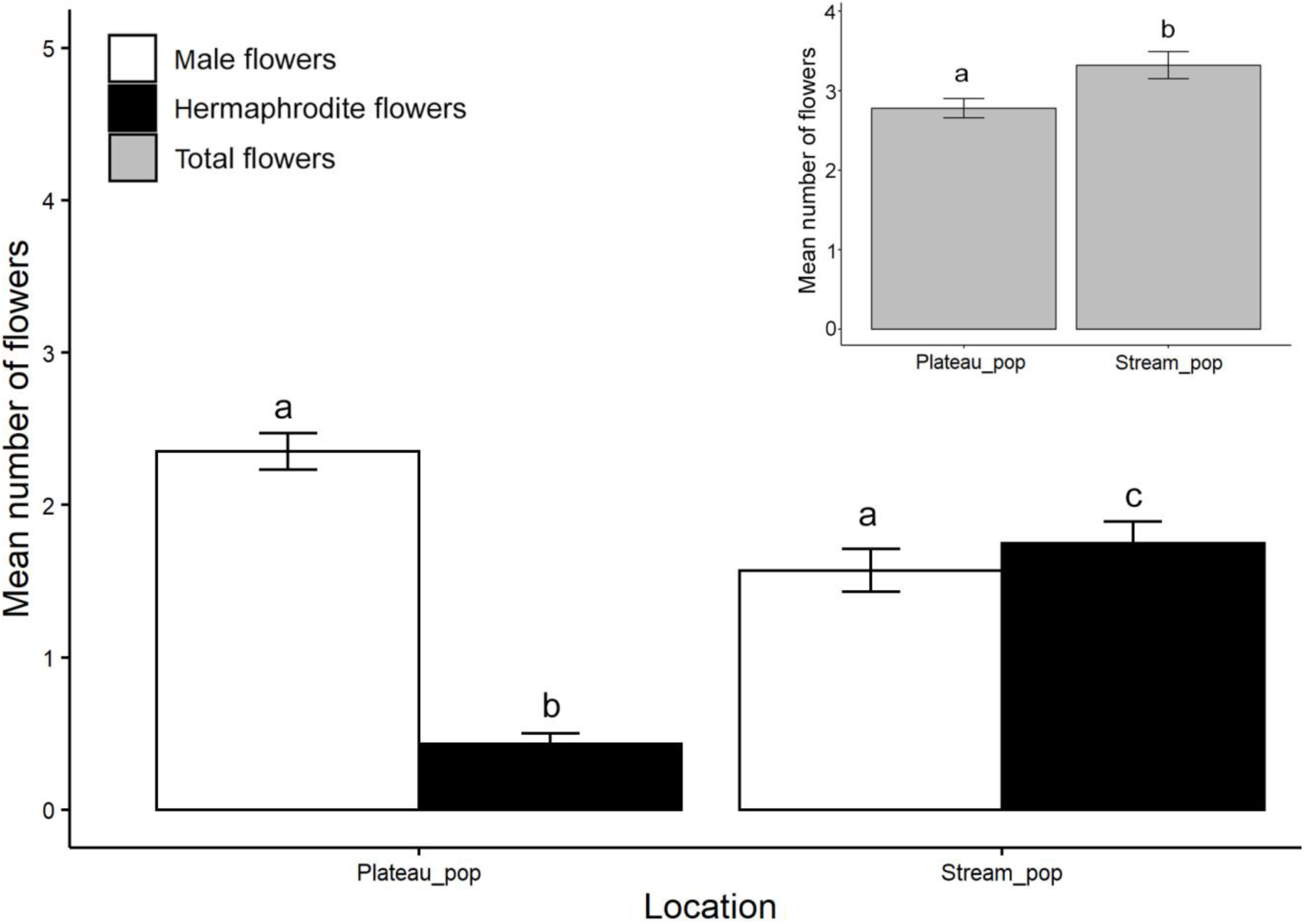
Natural floral sex distribution observed in *M. simplex* in 2019 between two populations (Mean±SE) (Kruskal-Wallis chi-squared = 123.53, df = 3, *p*-value <2.2^e-16^, N = 100 per population; Online Resource 1: Table 2; P<0.025) (Inset: Total number of flowers across the two populations; t = -2.5646, df = 176.65, *p*-value = 0.0116). Male flowers are represented by open squares, hermaphrodite flowers represented by filled squares and the gray squares represent total (male + hermaphrodite) flowers. Columns to be compared within population and within sexes across the two populations only.

Within an inflorescence, male and hermaphrodite flowers were present in all the three positions i.e. apical, middle and basal positions (Fig. 1c), and the mean number of male flowers varied across all these three positions during our four-week observation period. The number of male flowers were relatively consistent in the basal position throughout the plants’ flowering season than in the middle position or apical position (Fig. 2 b - e; Online Resource 1: Table 3, 4). We documented the highest number of male flowers in the middle position of the inflorescence, whereas the number of hermaphrodite flowers was consistent at all positions over the flowering season (Fig. 2 b - e; Online Resource 1: Table 3, 4).

The total number of flowers were significantly higher in the stream population than in the plateau population (Mean±SE: Plateau_pop = 2.78±0.12 vs. Stream_pop = 3.32±0.17; t = - 2.5646, df = 176.65, *p*-value = 0.0116; Fig. 3 inset) and the proportion of male:hermaphrodite flowers were similar in the stream population (Mean±SE: Stream_pop H = 1.75±0.14 vs. Stream_pop M = 1.57±0.14; *p*-value = 0.1614; Fig. 3). Number of male flowers was significantly higher in the plateau population over the stream population (Mean±SE: Plateau_pop M = 2.35±0.12 vs. Stream_pop M = 1.57±0.14; *p*-value = 0.0000; Fig. 3; Online Resource 1: Table 2).

The mean fruit set per plant showed no effect with the total number of male, hermaphrodite, or total number of flowers present per plant, respectively in the plateau_pop (hermaphrodite: *r* ^*2*^ = -0.0457, df = 18, *p*-value = 0.6852; Online Resource 1: Fig. 3a; male: *r* ^*2*^ = - 0.04999, df = 18, *p*-value = 0.761; Online Resource 1: Fig. 3b; total flowers: *r* ^*2*^ = -0.04254, df = 18, *p*-value = 0.6412; Online Resource 1: Fig. 3c), and in the stream_pop (hermaphrodite: *r* ^*2*^ = - 0.0.05543, df = 18, *p*-value = 0.9637; Online Resource 1: Fig. 3d; male: *r* ^*2*^ = 0.1125, df = 18, *p*-value = 0.08134; Online Resource 1: Fig. 3e; total flowers: *r* ^*2*^ = 0.04828, df = 18, *p*-value = 0.1781; Online Resource 1: Fig. 3f), respectively.

### Pollination biology

Flowers of both the sexes opened between 12:30 hrs. to 16:00 hrs., and their mean pollen count ranged from 5000-10000 pollen grains per anther and no significant difference in the pollen quantity between the two sexes was observed (t = -0.67913, df = 60, *p*-value = 0.4997; Azad 2018, Albal 2019, Nawge 2019). The major pollinators to *M. simplex*, in the order of visitation rates from highest to lowest were *Apis dorsata* (Mean number of visits per flower per hour±SE = 1.01±0.18), *Apis cerana* (0.86±0.10), *Apis florea* (0.76±0.22), and *Amegilla spp. (*Zonamegilla, 0.3±0.09) respectively. *Amegilla spp*. showed significantly lower visitation than all the other pollinators (Kruskal-Wallis chi-squared = 6.5702, df = 3, *p*-value = 0.09; *p*-values: *Amegilla spp*. vs *A. cerana* = 0.0074, *Amegilla spp*. vs *A. dorsata* = 0.0058, *Amegilla spp*. vs *A. florea* = 0.0469), whereas the visitation rates among *A. cerana, A. florea* and *A. dorsata* were not significantly different (*p*-values: *A. cerana* vs *A. dorsata* = 0.3822, *A. cerana* vs *A. florea* = 0.4278, *A. dorsata* vs *A. florea* = 0.3748; Azad 2018). Other pollinators observed were *Paragus spp*. (Hoverfly), *Lasioglossum spp*., *Pseudapis spp*. and one species that we could not identify which will henceforth be referred to as UnID sp. (Online Resource 1: Fig. 4). Mean pollinator visitation rates were observed to be high at the beginning of the anthesis (1.74-1.94 flowers from 13:00 hrs. to 15:00 hrs.) and it decreased with time (0.94-1.32 flowers from 15:00 hrs. till anthesis end; Azad 2018).

**Fig. 4.**
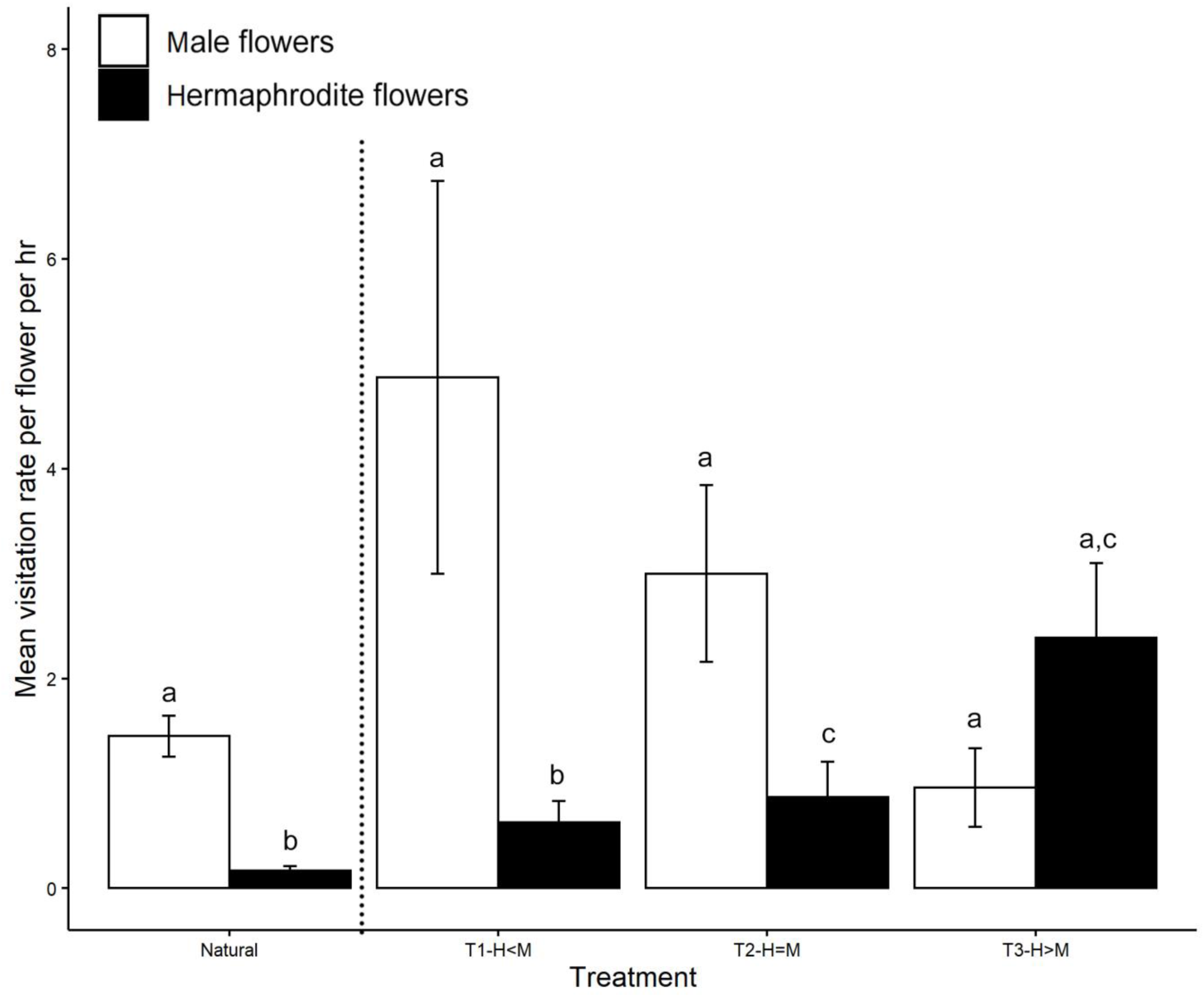
Mean pollinator visitation rate per flower per hour (Mean±SE) observed in the manipulative experiment where three treatments with varying numbers of hermaphrodite and male flowers were used to test pollinator preference towards a sex; (Kruskal-Wallis chi-squared = 63.4196, df = 7, *p*-value = 3.122^e-11^, N = 8 trials for each treatment; P<0.025). Male flowers are represented by open squares and hermaphrodite flowers are represented by the filled squares. The first two columns represent the natural visitation rate observed per flower per hour for both male and hermaphrodite, respectively. Different alphabets depict significantly different values (Online Resource 1: Table 5; P<0.025).

### Sex-specific pollinator preference experiment

In all the experimental setups in the manipulated choice experiment, pollinator visitations were highest to the most abundant sex (Kruskal-Wallis chi-squared = 65.3103, df = 7, *p*-value = 1.303^e-11^; Fig. 4; Online Resource 1: Table 5). Pollinator visitation rates were significantly different between hermaphrodite flowers in the unmanipulated, wild set up (control) and hermaphrodite flowers in all the experimental setups (Fig. 4; Online Resource 1: Table 5). However, the visitation rates to the male flowers in the control were not different from those to male flowers in all the experimental setup (Fig. 4; Online Resource 1: Table 5). Display size (total number of flowers per trial) had no effect on the visitation rates in any of the manipulated treatments (*r* ^*2*^ = -0.03765, df = 21, *p*-value = 0.6309; Online Resource 1: Fig. 5).

**Fig. 5.**
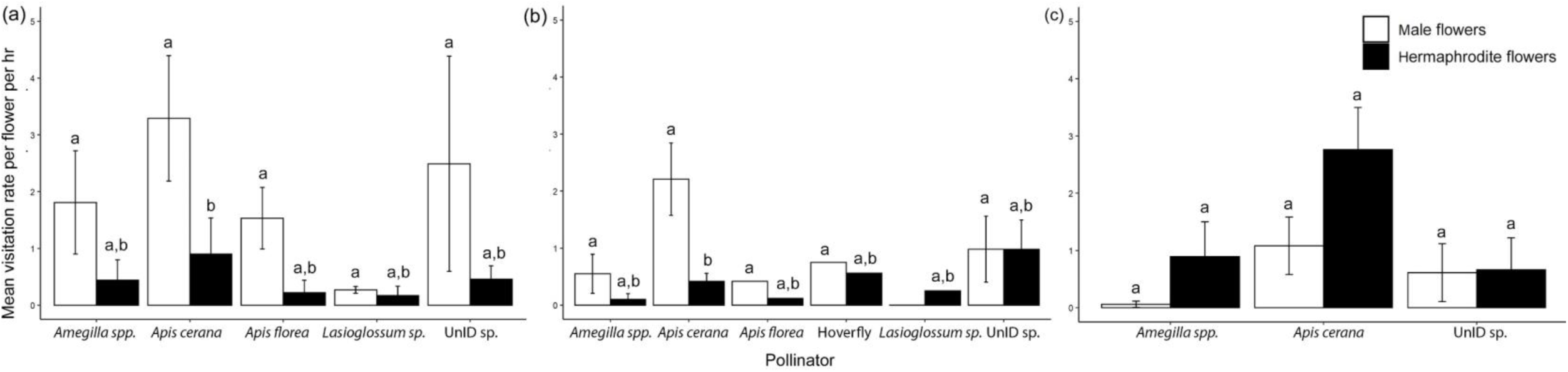
Pollinator-wise mean visitation rate per flower per hour in the manipulative experiment on *M. simplex* (Mean±SE) (a) Treatment 1-H < M (Kruskal-Wallis chi-squared = 13.8517, df = 9, *p*-value = 0.1277; P<0.025; Online Resource 1: Table 6), (b) Treatment 2-H = M (Kruskal-Wallis chi-squared = 15.605, df = 11, *p*-value = 0.1565; P<0.025; Online Resource 1: Table 6), (c) Treatment 3-H > M (Kruskal-Wallis chi-squared = 9.2848, df = 5, *p*-value = 0.09823; P<0.025; Online Resource 1: Table 6). Male flowers are represented by open squares and hermaphrodite flowers are represented by filled squares. Different alphabets depict significantly different values (Online Resource 1: Table 6; P<0.025).

Quantification of pollinator-wise visitation rates in the manipulated experiments show that visitations by *Apis cerana, Amegilla spp*. and UnID sp. were higher to male flowers in treatment 1 where male flowers were more abundant than hermaphrodite flowers (Mean visitation rate per flower per hr±SE:-H < M: *A. cerana* = 3.29±1.11, *Amegilla spp*. = 1.81±0.91, UnID sp. = 2.49±1.89 flowers; Kruskal-Wallis chi-squared = 13.8517, df = 9, *p*-value = 0.1277; Fig. 5a; Online Resource 1: Table 6). In treatment 3, *Apis cerana, Amegilla spp*., and UnID sp. showed high visitation to hermaphrodite flowers (H > M: *A. cerana* = 2.76±0.74, *Amegilla spp*. = 0.89±0.61, UnID sp. = 0.66±0.56 flowers; Kruskal-Wallis chi-squared = 9.2848, df = 5, *p*-value = 0.09823; Fig. 5c; Online Resource 1: Table 6). We observed visitation by hoverflies only in treatment 2 (H = M: Visitation rate per hermaphrodite flower = 0.56, visitation rate per male flower = 0.75; Kruskal-Wallis chi-squared = 15.605, df = 11, *p*-value = 0.1565; Fig. 5b; Online Resource 1: Table 6).

## Discussion

### Population-level sex ratios

The optimal allocation hypothesis proposes that between a costly and a cheaper sex, in a resource-limited environment, the sex which requires the least resources will be preferred. From a plant’s perspective, the male flowers are considered a cheaper sex to produce because there is no investment towards fruit development, unlike the hermaphrodite flowers which produces both viable pollen and ovules (Bertin 1982a, Anderson and Symon 1989, Kaul et al. 2002, Narbona et al. 2002, Obeso 2002, Verdú et al. 2007, Zhang and Tan 2009). Here, we show that the *M. simplex* population on Kaas plateau has a higher density of male flowers and a slight reduction in the number of male flowers and an increase in hermaphrodite flowers were observed only among the individuals present along the stream. We consider the stream population of *M. simplex* to be growing in better resource conditions than the plateau population because of the presence of water, possible higher nutrient availability due to dissolved/associated nutrients in the water, and deeper soil substrate for the plants (Thorpe et al. 2018). This suggests that although the general tendency of *M. simplex* is to produce more male flowers over hermaphrodites, water and water-dissolved minerals may be major limiting resources and in the absence of resource constraints the plants can produce not only more flowers but also higher number of hermaphrodite flowers.

### Temporal shift in floral sex

Resources available and utilized by plants often fluctuate with time and age of the plant (Stephenson 1982, Solomon 1985). We observed an initial increase in the number of flowers followed by a decreasing trend in the total number of flowers being produced per plant throughout its flowering period (Fig. 2a inset). Within this trend, male flowers were always produced in higher proportions than the hermaphrodite flowers and interestingly, in the end of the flowering season (i.e. fourth week) we observed a reduction in the number of male flowers instead of the number of hermaphrodite flowers (Fig. 2a). Based on these observations we conclude that although resource constraints negatively affect the total production of flowers, the reduction in flowers was heavily biased against male flowers, while the number of hermaphroditic flowers did not decrease with time in *M. simplex*.. Contrary to our observation, it has been shown that under declining resource levels plants may opt to increase their fitness by producing the relatively inexpensive male flowers over the expensive female flowers (May and Eugene Spears Jr 1988). However, in *C. benghalensis* it has been reported that a plant will opt to drop a flower rather than drop a fruit, under resource limiting conditions, because the nutrient resources that would get committed to fruits that are aborted would be saved halfway (Kaul and Koul 2008). We propose that a similar strategy may be operational in *M. simplex* wherein under availability of resources *M. simplex* produces an excess of male flowers, which are the first ones to be sacrificed when the plants face resource limitation as the flowering season progresses (Fig. 2a). The lack of change in the number of hermaphrodite flowers in different stages of plant’s flowering phenology also suggests that the hermaphrodite flowers may already be produced at an optimal number, and hence their numbers do not decline despite declining resources (Fig. 2a).

Distribution of the two sexes within an inflorescence has been shown to be dependent on resource allocation (Diggle 1997, Miller and Diggle 2003), and it has been observed that the differential positioning of the sexes within an inflorescence can affect outcrossing rates (Orth and Waddington 1997, Lin and Forrest 2019). Studies in *Solanum* (Diggle 1994, 1997, Miller and Diggle 2003), *Caesalpinia* (Lewis and Gibbs 1999), *F. virginiana* (Ashman and Hitchens 2000) and in other Commelinaceae genera (Kaul et al. 2002, Kaul and Koul 2008) have shown that higher accessibility to nutrient resources at the base of an inflorescence (than at the distal end) can result in the presence of higher resource-demanding hermaphrodite flowers in the basal position while males are relegated to the middle and apical positions of an inflorescence. We found a similar pattern in *M. simplex*, where a greater number of male flowers were produced at the middle and the apical positions on an inflorescence. However, unlike other plants where hermaphrodites exclusively occupy the basal positions in an inflorescence, in *M. simplex* hermaphrodites were also present in the apical position. In fact, the distribution of hermaphrodite flowers was not significantly different across all the three positions, over the course of the plant’s flowering season. This implies that any manipulation in display size is carried out by increasing or decreasing the male flowers thus regulating the floral display at a much lesser cost (as proposed in *Solanum* by Anderson and Symon 1989).

### Pollination biology

Pollinators have been shown to display higher preference to larger inflorescences, ultimately resulting in higher reproductive success (Stephenson 1981, Harder and Barrett 1995, Ohashi and Yahara 2001). Hence the disproportionate distribution of male and hermaphrodite flowers within the inflorescence of an andromonoecious species may be a result of indirect pollinator selection on inflorescence size: that is larger inflorescences are better at attracting pollinators but under nutrient resource limitation, this can be achieved only by producing more of the cheaper/economical male flowers (Sandring and Ågren 2009). Thus our results support the pollinator attractor hypothesis as proposed by Zhang and Tan (2009) and sex allocation theory as proposed by Elle and Meagher (2000) where they predicted that male flowers can increase the floral display and their attractiveness to pollinators at a lower cost through male function than female function. The comparable fruit set values between the plateau and the stream population of *M. simplex* also suggests that reproductive costs were not compromised while decreasing male flowers due to resource constraints in the plateau population.

The dominant pollinator assemblage visiting *M. simplex* consisted mainly of different species of bees from the two genera common in the Indian tropics - *Apis* and *Amegilla*. All pollinators that visit *M. simplex* are generalists and are known to be common pollinators in grassland habitats (Corlett 2011, Dhargalkar 2019, Nawge 2019). There are at least 68 plants co-flowering with *M. simplex* on the Kaas plateau (Dhargalkar 2019) and parallel studies on the pollination of plants in the Kaas community showed that both *Apis cerana* and *Apis florea* switch to *M. simplex* from *M. lanuginosa* (anthesis time 10.30 hrs.-13.00 hrs.) and various *Impatiens* species as soon as the flowers of *M. simplex* open i.e. at 12:30 hrs. up to 16:00 hrs. (Albal 2019). This suggests that although the pollinators are generalists, these pollinators may be displaying temporal specialization towards *M. simplex*. Future studies will focus on temporal specialization of the bee communities on Kaas plateau, which will provide insight on pollinator preference towards *M. simplex* in a multi-species environment.

### Sex-specific pollinator preference

Several studies have shown that pollinators avoid functionally female flowers because of decreased pollen or absence of pollen (Biezychudek 1987, Eckhart 1991), floral size (Ashman et al. 2000), floral scents (Ashman et al. 2005) and pollen-pistil interference (Solomon 1985, Elle and Meagher 2000). However, in *M. simplex* both male and hermaphrodite flowers showed comparable pollen production. This has been shown in other andromonoecious systems as well such as in *Solanum carolinense* (Solomon 1986), *Olea europaea* (Cuevas and Polito 2004) and *Dichorisandra spp*. (Sigrist and Sazima 2015). As previously discussed, we observed that nutrient resources affected the number of male flowers and not the number of hermaphrodite flowers, and the rate of fruit sets were observed to be similar between the plateau and the stream populations (Mean±SE: Plateau_pop = 3.22±0.23 vs. Stream_pop = 3.55±0.21). This suggests that reproductive success is limited by the availability of hermaphrodite flowers and not pollinator visitations. The lack of preference for either of the sexes by the pollinators on Kaas plateau is further supported by our observations from the manipulative experiment where all pollinators except *A. cerana* displayed lack of preference towards either of the floral sexes. Further, this also fails to support the hypothesis that male-dominant inflorescences of *M. simplex* may be under pollinator-mediated selection due to direct pollinator preference for any sex (see Solomon 1985, Eckhart 1991, Elle and Meagher 2000). However, since pollinator visitations were observed to be density dependent (i.e. floral density, compare treatment 3 with treatment 1, Fig. 5c) it is possible that there is an indirect pollinator-mediated selection on male flowers through preference for larger inflorescence sizes.

Our understanding of pollinator services by bees primarily come from *Bombus* species (Chittka et al. 1997) and *Apis mellifera* (Martin 2004) and studies on other bees from India are very rare (see Somanathan and Borges 2001). At least five species of bees visited *M. simplex* and in the choice experiment, we observed inter-species behavioral differences among bees both in their floral preferences and visitation patterns, respectively. Almost all bees displayed higher visitation to any sex which was present in greater density, suggesting that pollinator visitations are predominantly sex-nonspecific and density-dependent. A slight preference for sex was only observed in *A. cerana* and *Amegilla* in treatment 2 (H = M) where male flowers were preferred over hermaphrodite flowers (Fig. 5b). Bees are known for both their short-term and long-term memory retention and they can show learning through foraging experience (Cresswell and Galen 1991, Schiestl and Johnson 2013) and we propose that the higher visitation to male flowers in treatment 2 (Fig. 5b) may be a result of short-term memories and habituation in bees since in natural conditions bees may encounter more male flowers than hermaphrodites.

We conclude that in *M. simplex* environmentally driven resource constraints play a bigger role than pollinator preference towards a specific sex in determining floral sex expression. However, we cannot dismiss an indirect role of pollinator-driven constraints which may occur through selection on the display size as proposed in the pollinator attractor hypothesis (Zhang and Tan 2009). In a resource-limited environment such as the one observed at Kaas plateau, *M. simplex* can thus increase its display size by producing the less expensive male flowers over hermaphrodite flowers.

Energy constraints will have a direct cost on floral production if the two sexes require differential resource allocation. As a result, it can be expected that the sex of the flowers will be selected accordingly. Using the *M. simplex* system, we show that floral sex expression is directly influenced by the availability of nutrient resources in the plants’ environment and indirectly affected by the pollinators’ preference for larger display sizes of the inflorescence. Thus, a combination of both environmental factors as well as ecological factors may be responsible for the evolution and maintenance of andromonoecy in mass-flowering plants such as *M. simplex* in the tropics.

## Supporting information

Online Resource 1

## Acknowledgements

The authors thank the Ministry of Human Resource Development, India (MHRD) and the National Geographic Society for research funds to VG, IISER Bhopal for the infrastructure, DST for INSPIRE fellowship to AA, AG, and SS. We thank the PCCF of Maharashtra Forest Department, DFO of Satara Division (Forest Department) and the Joint Forest Management Committee (JFMC), Kaas, Satara for permitting our fieldwork. We also thank Dr. Natapot Warrit (Chulalongkorn University, Thailand) and Dr. Ximo Mengual (Research Museum Alexander Koenig, Germany) for their help with taxonomic identification of the pollinators. Finally, we thank all fellow TrEE lab members for their advice, discussion, and support.

## Compliance with ethical standards

### Conflict of interest

Authors have declared no conflicts of interest.

### Statement of human and animal rights

This article does not contain any studies with human participants or animals performed by any of the authors.

## Notes

### Competing Interest Statement

The authors have declared no competing interest.

