## Supplementary material for "When to be a male? Role of resource-limitation and pollinators in determining floral sex in an andromonoecious spiderwort": Online Resource 1

### Electronic supplementary material

In 2017, the mean fruit set per plant was not affected by the mean number of hermaphrodite flowers (Online Resource 1: Fig. 6a: $r^2$ = -0.02284, df = 24, *p*-value = 0.5126) or mean number of male flowers (Online Resource 1: Fig. 6b: $r^2$ = 0.0405 , df = 24, *p*-value = 0.1646) respectively, but showed a positive effect with the total number of flowers present per plant (Online Resource 1: Fig. 6c: $r^2$ = 0.12, df = 24, *p*-value = 0.04642). However in 2019, the mean fruitset per plant showed no effect with the mean number of hermaphrodite flowers (Online Resource 1: Fig. 6d: $r^2$ = -0.01361, df = 38, *p*-value = 0.4943), mean number of male flowers (Online Resource 1: Fig. 6e: $r^2$ = 0.05935 , df = 38, *p*-value = 0.07059), or total number of flowers present per plant, respectively (Online Resource 1: Fig. 6f: $r^2$ = -0.002529, df = 38, *p*-value = 0.3484).

The 2017 and 2018 census data were pooled to observe the overall trend in the gender distribution of the population of *M. simplex* (Online Resource 1: Fig. 7). The number of male flowers remained consistently higher than the hermaphrodite flowers in both years, where as the number of hermaphrodite flowers itself remained similar in both years, supporting our results.

### Tables

**Table 1** Kruskal-Wallis test results of floral sex distribution (M = male, H = hermaphrodite, T = total flowers) in *M. simplex* across four weeks in 2018 showing the temporal variation in sex distribution (Fig. 2a; Kruskal-Wallis chi-squared = 38.7067, df = 7, *p*-value= 0.00; Fig 2a inset; Kruskal-Wallis chi-squared = 3.2847, df = 3, *p*-value = 0.3498; P<0.025).

| Week | Week | 1 | | | 2 | | | 3 | | | 4 |
|---|---|---|---|---|---|---|---|---|---|---|---|
| | Sex/total | H | M | T | H | M | T | H | M | T | H |
| 1 | M | **0.0154*** | | | | | | | | | |
| 2 | H | 0.2843 | - | - | | | | | | | |
| | M | - | 0.1181 | - | **0.0000*** | | | | | | |
| | T | - | - | 0.4066 | - | - | | | | | |
| 3 | H | 0.2218 | - | - | 0.4222 | - | - | | | | |
| | M | - | 0.1549 | - | - | 0.4328 | - | **0.0000*** | | | |
| | T | - | - | 0.2566 | - | - | 0.3382 | - | - | | |
| 4 | H | 0.4669 | - | - | 0.3131 | - | - | 0.2472 | - | - | |
| | M | - | 0.1762 | - | - | **0.0172*** | - | - | 0.0258 | - | 0.0947 |
| | T | - | - | 0.1399 | - | - | 0.0939 | - | - | 0.0414 | - |

**Table 2** Kruskal-Wallis test results of floral sex distribution (M = male, H = hermaphrodite) in *M. simplex* between two populations (Fig. 3; Kruskal-Wallis chi-squared = 123.5274, df = 3, *p*-value = 0; P<0.025).

| Comparative groups | *p*-value |
|---|---|
| Plateau_pop M: Plateau_pop H | **0.0000*** |
| Stream_pop H: Plateau_pop H | **0.0000*** |
| Stream_pop M: Plateau_pop M | **0.0000*** |
| Stream_pop M: Stream_pop H | 0.1614 |

**Table 3** Kruskal-Wallis test results of temporal variation in floral sex distribution (M- male, H- hermaphrodite) within an inflorescence in *M. simplex* (Fig. 2b-e; P<0.025). Floral sex was quantified at three positions within an inflorescence: basal, middle (mid) and apical (see Fig. 1c).

| Comparative groups | Week 1 | Week 2 | Week 3 | Week 4 |
|---|---|---|---|---|
| | *p*-value | *p*-value | *p*-value | *p*-value |
| Basal H: Basal M | **0.0014*** | 0.0252 | 0.0377 | 0.3547 |
| Basal H: Mid H | 0.3912 | 0.3744 | 0.2759 | 0.1844 |
| Basal M: Mid M | **0.0216*** | 0.0373 | 0.0377 | **0.0027*** |
| Mid H: Mid M | 0.2472 | **0.0003*** | **0.0015*** | 0.0661 |
| Basal H: Apical H | 0.0590 | 0.5000 | 0.4543 | 0.4132 |
| Mid H: Apical H | 0.0329 | 0.3744 | 0.3155 | 0.1318 |
| Basal M: Apical M | **0.0000*** | 0.2817 | 0.4686 | 0.0700 |
| Mid M: Apical M | **0.0058*** | **0.0091*** | 0.0316 | 0.0967 |
| Apical H: Apical M | 0.5000 | 0.0840 | 0.0565 | 0.0930 |

**Table 4** Mean±SE for the distribution of floral sex (M- male, H- hermaphrodite) at three positions- basal, middle, and apical- within an inflorescence in *M. simplex* (Fig. 2b-e).

| Position | Week | Week 1 | Week 2 | Week 3 | Week 4 |
|---|---|---|---|---|---|
| | Sex | | | | |
| Basal | M | 0 | 1±0.39 | 1.1±0.48 | 0.8±0.33 |
| | H | 0 | 0.3±0.21 | 0.2±0.13 | 0.3±0.21 |
| Middle | M | 1.2±0.36 | 2.7±0.6 | 3.1±0.74 | 1.6±0.48 |
| | H | 1.1±0.53 | 0.4±0.22 | 1±0.73 | 0.6±0.22 |
| Apical | M | 2.8±0.59 | 1.1±0.31 | 1±0.3 | 0.2±0.13 |
| | H | 0.7±0.3 | 0.3±0.21 | 0.3±0.3 | 0.3±0.15 |

**Table 5** Kruskal-Wallis test results of mean pollinator visitation rate per flower per hour from the manipulative experiments in *M. simplex* (Fig. 6; Kruskal-Wallis test chi-squared= 63.4196, df=7, *p*-value=3.122$^{e-11}$; P<0.025).

| Treatment | | Natural | | T1 | | T2 | | T3 |
|---|---|---|---|---|---|---|---|---|
| | Sex | H | M | H | M | H | M | H |
| Natural | M | **0.0000*** | | | | | | |
| T1 | H | 0.0545 | | | | | | |
| | M | | 0.0412 | **0.0060*** | | | | |
| T2 | H | **0.0106*** | | 0.3611 | | | | |
| | M | | 0.1049 | | 0.3053 | **0.0248*** | | |
| T3 | H | **0.0000*** | | 0.0277 | | 0.0457 | | |
| | M | | 0.2014 | | **0.0235*** | | 0.0544 | 0.0884 |

**Table 6** Kruskal-Wallis test results for pollinator-wise mean visitation rate per flower per hour in the manipulative experiments for *M. simplex* (Fig. 7; P<0.025) (M- male, H- hermaphrodite).

| Comparative groups | | *Amegilla spp.* H | *Amegilla spp.* M | *A. florea* H | *A. florea* M | *A. cerana* H | *A. cerana* M | UnID sp 3 H | UnID sp 3 M | UnID sp 4 H | UnID sp 4 M | Hoverfly H |
|---|---|---|---|---|---|---|---|---|---|---|---|---|
| *Amegilla spp.*M | H<M | 0.0912 | | | | | | | | | | |
| | H=M | 0.1086 | | | | | | | | | | |
| | H>M | 0.0795 | | | | | | | | | | |
| *A. florea* H | H<M | 0.3952 | - | | | | | | | | | |
| | H=M | 0.4422 | - | | | | | | | | | |
| | H>M | - | - | | | | | | | | | |
| *A. florea* M | H<M | - | 0.3616 | 0.0414 | | | | | | | | |
| | H=M | - | 0.4710 | 0.2966 | | | | | | | | |
| | H>M | - | - | - | | | | | | | | |
| *A. cerana* H | H<M | 0.3398 | - | 0.2533 | - | | | | | | | |
| | H=M | 0.1175 | - | 0.2744 | - | | | | | | | |
| | H>M | 0.1509 | - | - | - | | | | | | | |
| *A. cerana* M | H<M | - | 0.1650 | - | 0.3063 | **0.0119*** | | | | | | |
| | H=M | - | 0.0606 | - | 0.1426 | **0.0061*** | | | | | | |
| | H>M | - | 0.0663 | - | - | 0.0544 | | | | | | |
| UnID sp 3 H | H<M | 0.4181 | - | 0.4896 | - | 0.2926 | - | | | | | - |
| | H=M | 0.3419 | - | 0.4154 | - | 0.3765 | - | | | | | 0.2724 |
| | H>M | - | - | - | - | - | - | | | | | - |
| UnID sp 3 M | H<M | - | 0.1743 | - | 0.1179 | - | 0.0389 | 0.3781 | | | | - |
| | H=M | - | - | - | - | - | - | - | | | | - |
| | H>M | - | - | - | - | - | - | - | | | | - |
| UnID sp 4 H | H<M | 0.3365 | - | 0.2600 | - | 0.4686 | - | 0.2914 | - | | | - |
| | H=M | 0.0385 | - | 0.1346 | - | 0.1722 | - | 0.1995 | - | | | 0.4595 |
| | H>M | 0.3363 | - | - | - | 0.0621 | - | - | - | | | - |
| UnID sp 4 M | H<M | - | 0.4338 | - | 0.4304 | - | 0.2392 | - | 0.1518 | 0.1798 | | - |
| | H=M | - | 0.3711 | - | 0.3800 | - | 0.1244 | - | - | 0.4185 | | - |
| | H>M | - | 0.2708 | - | - | - | 0.2193 | - | - | 0.3536 | | - |
| Hoverfly H | H<M | - | - | - | - | - | - | - | - | - | - | - |
| | H=M | 0.1253 | - | 0.2063 | - | 0.3111 | - | - | - | - | - | - |
| | H>M | - | - | - | - | - | - | - | - | - | - | - |
| Hoverfly M | H<M | - | - | - | - | - | - | - | - | - | - | - |
| | H=M | - | 0.3105 | - | 0.3217 | - | 0.3259 | - | - | - | 0.3968 | 0.4293 |
| | H>M | - | - | - | - | - | - | - | - | - | - | - |

### Figures

**Fig. 1** Measurement of (a) floral width of male and hermaphrodite *M. simplex* flower, (b) 15 pairs arranged on a laminated graph sheet to observe sexual dimorphism (Photo courtesy: Azad G).

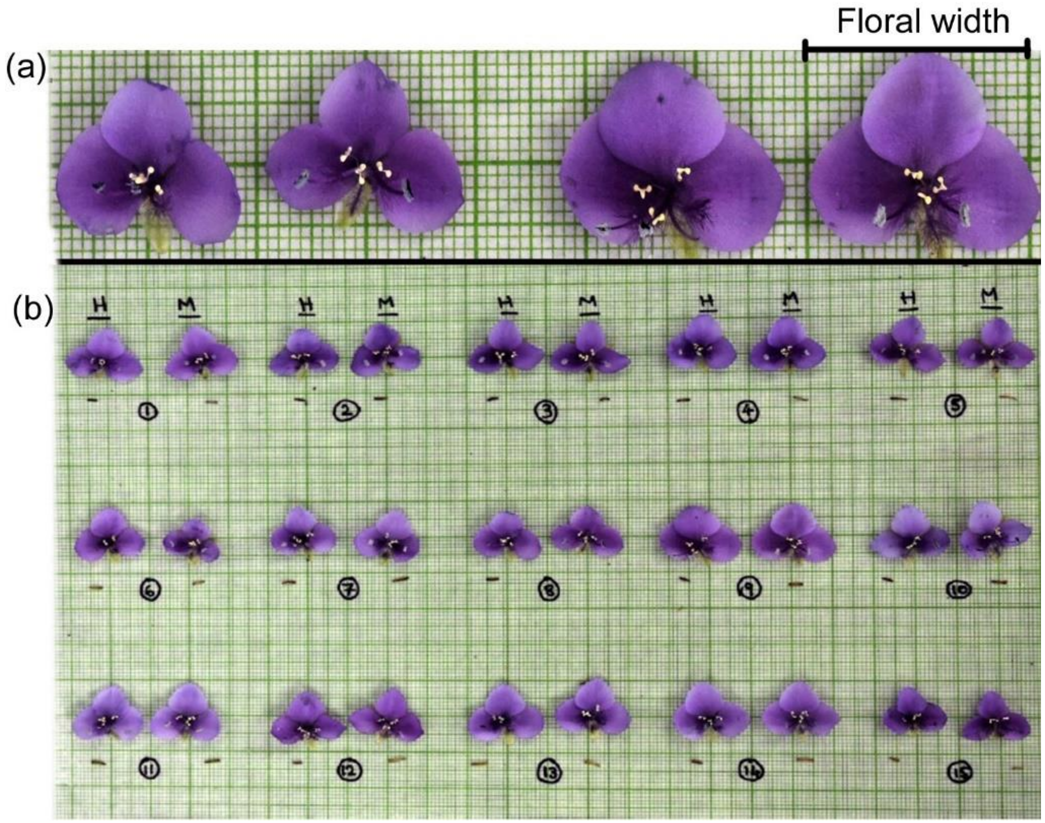

**Fig. 2** Manipulative choice experiment setup: (a) Treatment 1: H < M, (b) Treatment 2: H = M, (c) Treatment 3: H > M; (d) Experimental setup, 8-20 flowers in total; N = 8 trials. Illustration: Asawari Albal

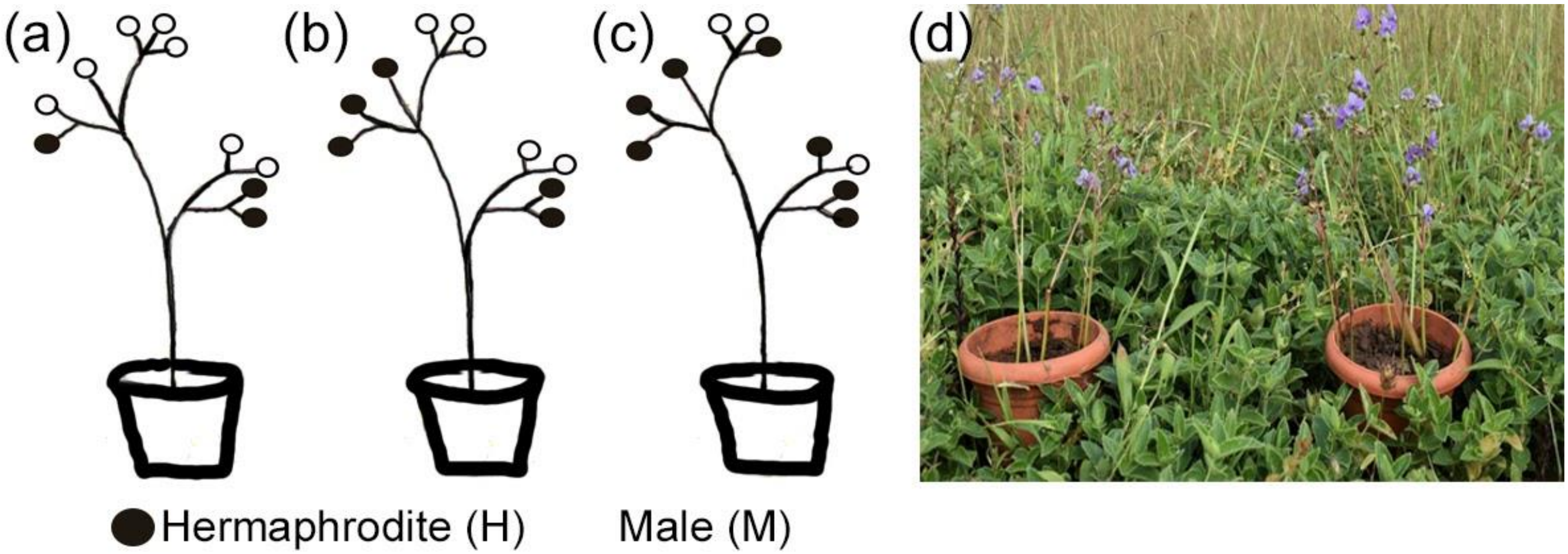

**Fig. 3** For plateau_pop in 2019, effect on mean fruitset by (a) mean number of male flowers ($r^2$ = -0.04999, df = 18, *p*-value = 0.761, N = 20), (b) mean number of hermaphrodite flowers ($r^2$ = -0.0457, df = 18, *p*-value = 0.6852, N = 20) and (c) mean number of flowers per plant ($r^2$ = -0.04254, df = 18, *p*-value = 0.6412, N = 20). For stream_pop in 2019, effect on mean fruitset by (f) mean number of male flowers ($r^2$ = 0.1125, df = 18, *p*-value = 0.08134, N = 20), (e) mean number of hermaphrodite flowers ($r^2$ = -0.05543, df = 18, *p*-value = 0.9637, N = 20) and (f) mean number of flowers per plant ($r^2$ = 0.04828, df = 18, *p*-value = 0.1781, N = 20).

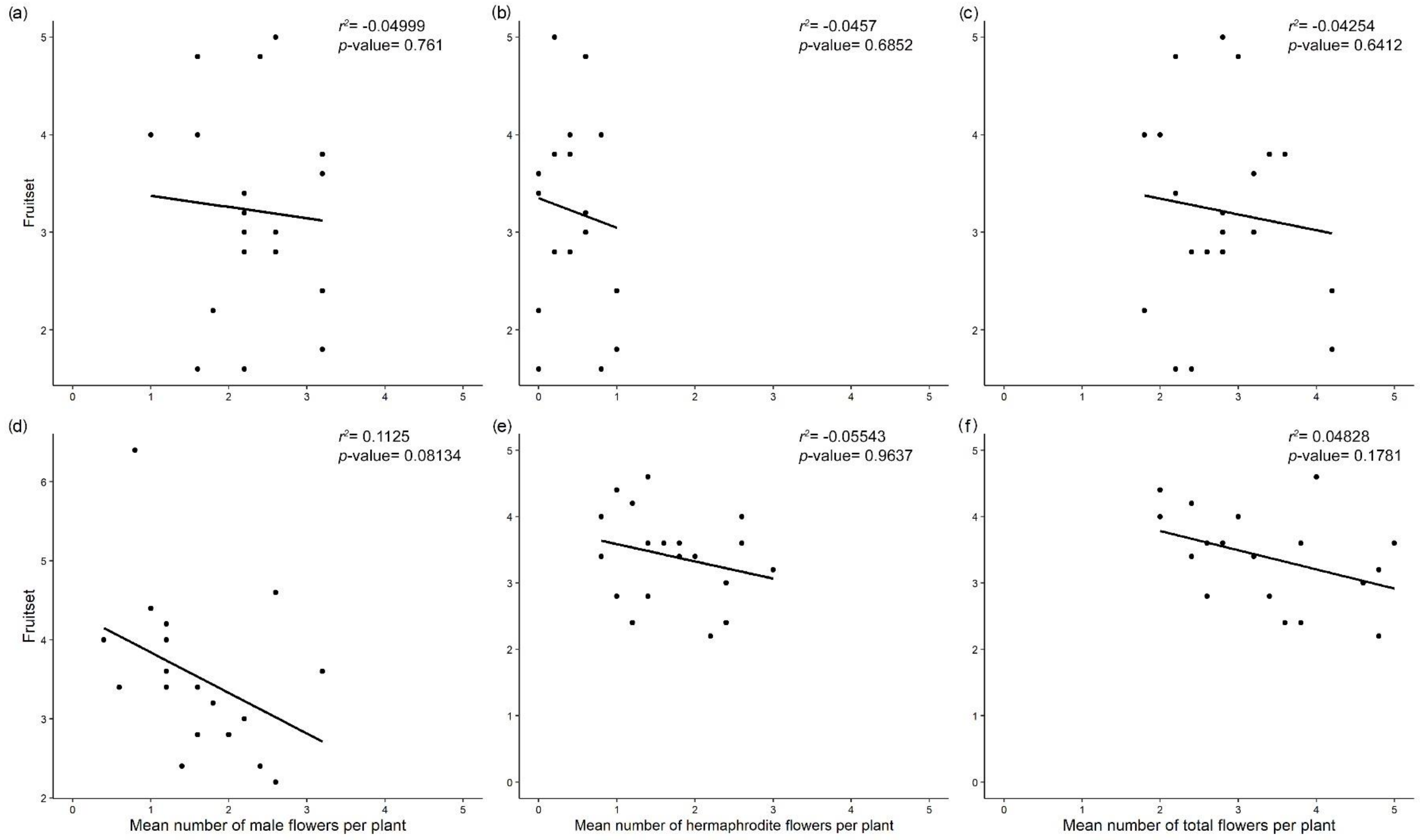

**Fig. 4** Pollinators of *M. simplex* from Kaas plateau: (a) and (b) *Amegilla spp. (Zonamegilla)*; (c) *Apis florea*; (d) *Apis cerana*; (e) *Apis dorsata*; (f) *Paragus sp.* (Hoverfly); (g) *Lasioglossum sp.*; (h) UnId sp. and (i) *Pseudapis sp.* Photo courtesy- (a) - (d), (f), (i) : Asawari Albal and (e), (g), (h): Saket Shrotri.

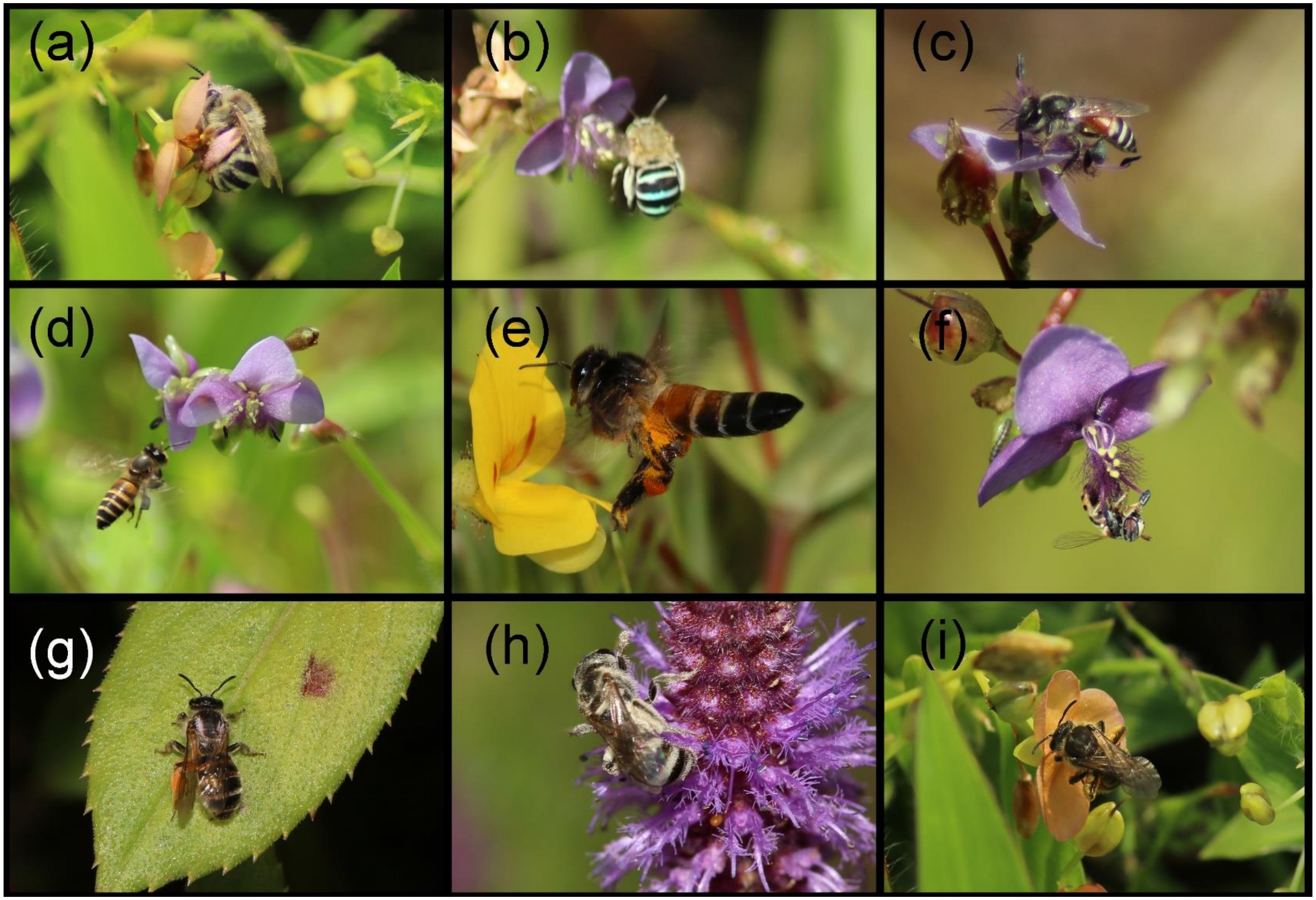

**Fig. 5** Effect of the total number of flowers on display and the mean visitation rate per hour ($r^2$ = -0.03245, df = 21, *p*-value = 0.5845, N = 22) in 2018. The filled circles with solid line represent treatment 1: H < M, filled triangles with dotted line represent treatment 2: H = M and filled squares with dash-dotted line represent treatment 3: H > M for the choice experiment.

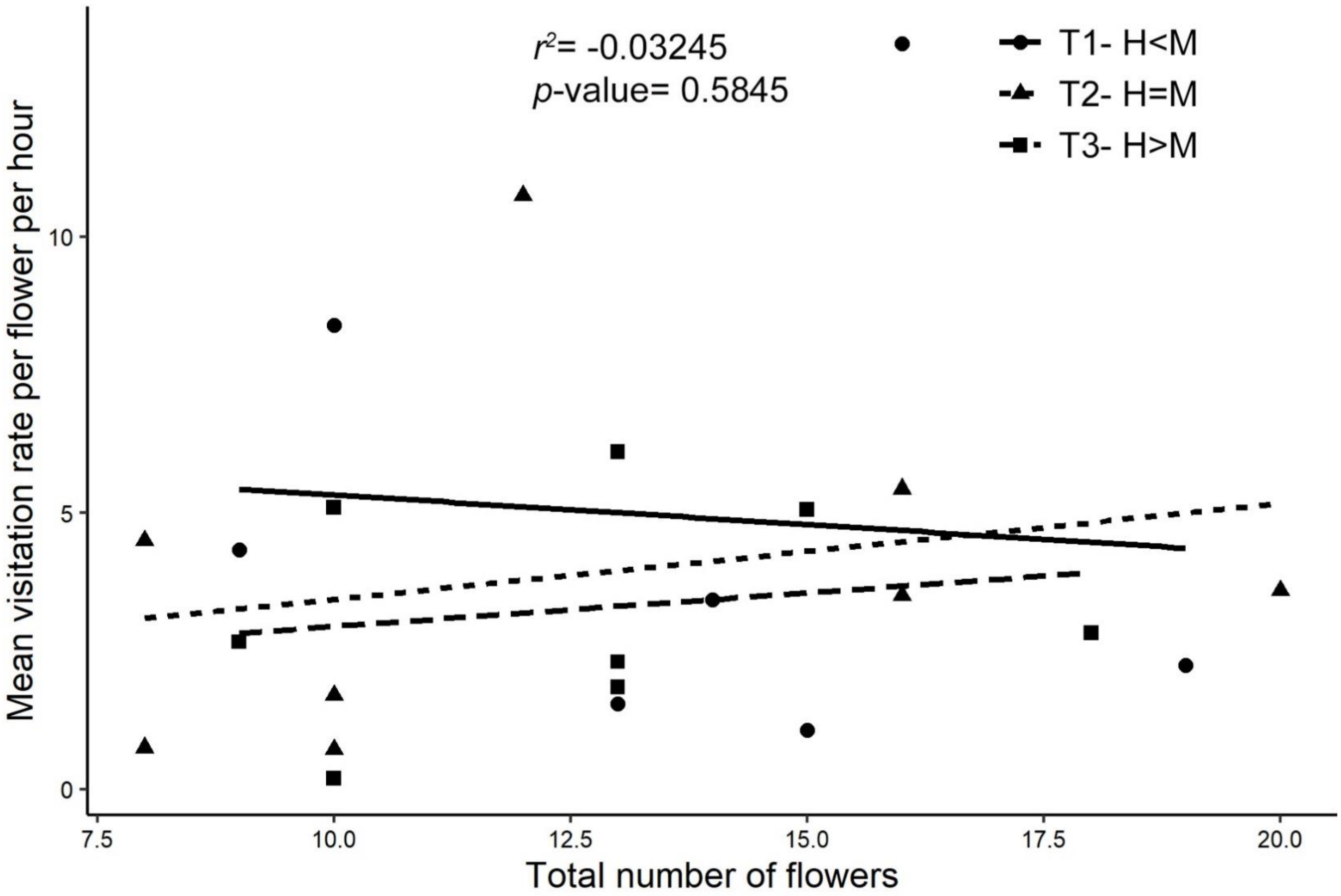

**Fig. 6** Effect on mean fruitset in 2017- (a) by mean number of male flowers ($r^2$ = 0.0405 , df = 24, *p*-value = 0.1646, N = 26), (b) by mean number of hermaphrodite flowers ($r^2$ = -0.02284, df = 24, *p*-value = 0.5126, N = 26) and (c) by mean number of total flowers per plant in ($r^2$ = 0.12, df = 24, *p*-value = 0.04642, N = 26). Effect of mean fruitset in 2019- (d) by mean number of male flowers ($r^2$ = -0.01361, df = 38, *p*-value = 0.4943, N = 40), (e) by mean number of hermaphrodite flowers ($r^2$ = 0.05935, df = 38, *p*-value = 0.07059, N = 40), (f) by mean number of total flowers ($r^2$ = -0.002529, df = 38, *p*-value = 0.3484, N = 40).

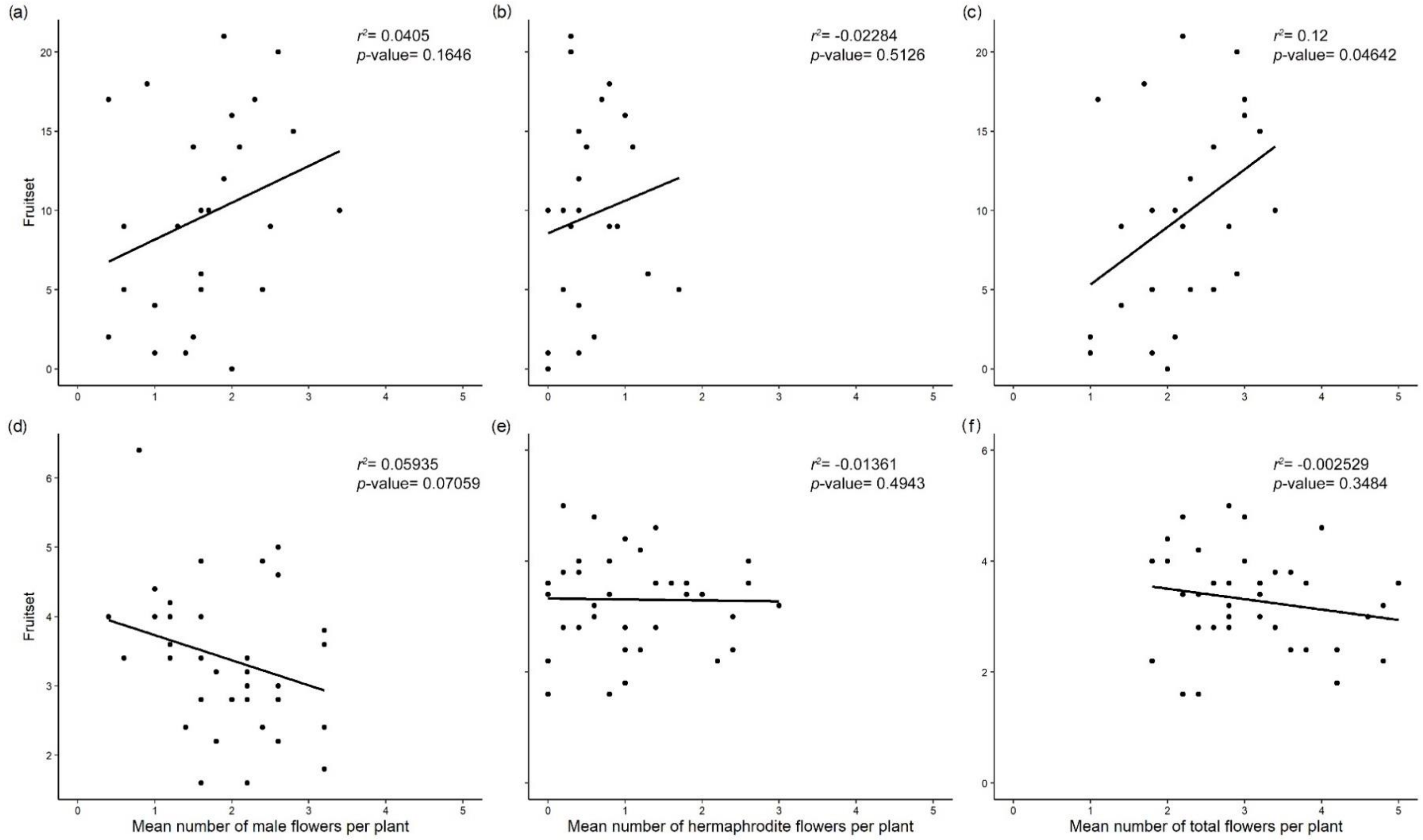

**Fig. 7** Variation in population-level sex ratios in *M. simplex* across two years 2017 and 2018 (Pooled data; Kruskal-Wallis chi-squared = 281.61, df = 7, $p$-value $<2.2^{e-16}$). Columns to be compared within the week and within sexes across all 4 weeks (Mean±SE). Significant $p$-values depicted by different alphabets.

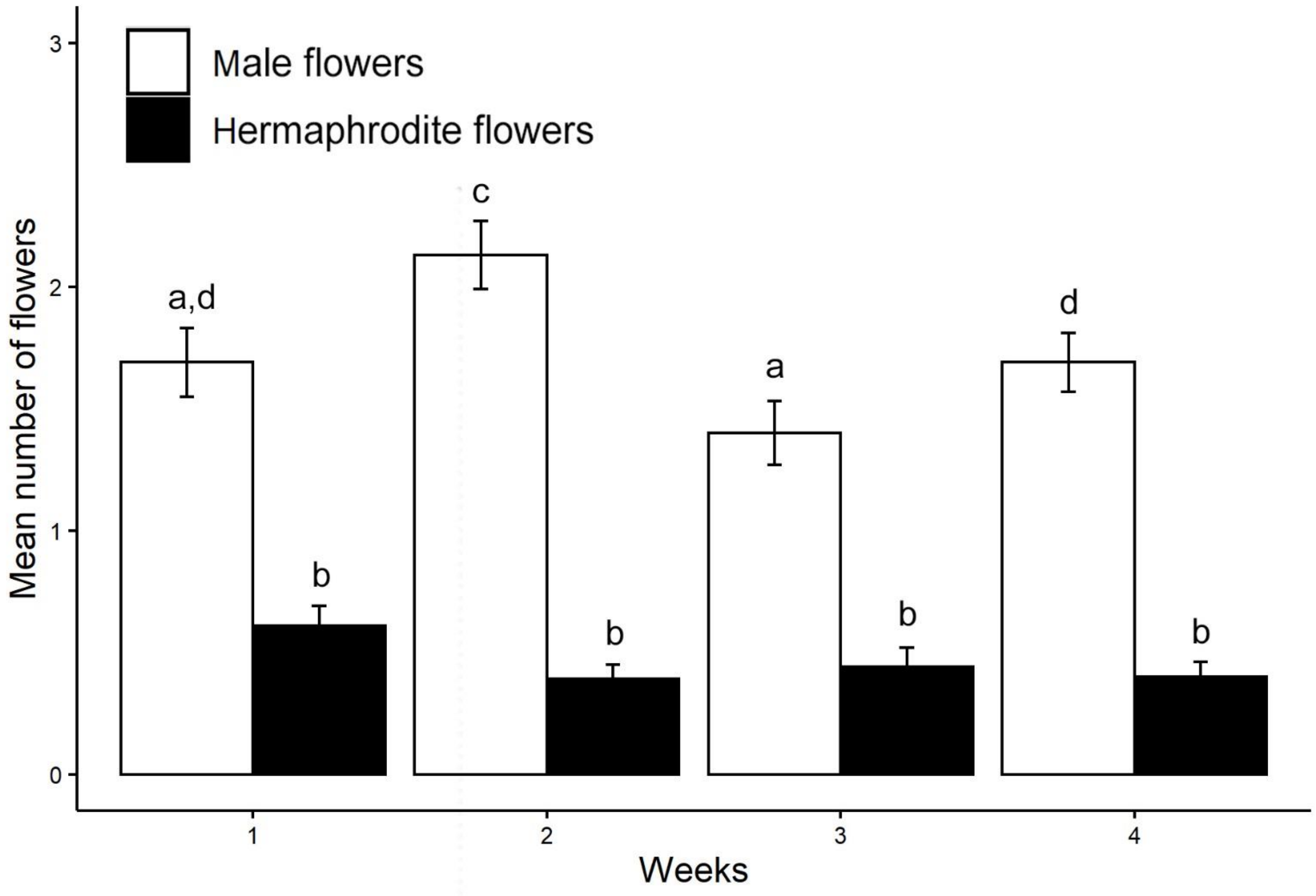
